# Putative staphylococcal enterotoxin possesses two common structural motifs for MHC-II binding

**DOI:** 10.1101/2023.07.14.548958

**Authors:** Shakilur Rahman, Saradindu Saha, Somdeb Bose Dasgupta, Amit Kumar Das

**Affiliations:** Department of Biotechnology, Indian Institute of Technology Kharagpur, Kharagpur 721302, West Bengal, India

**Keywords:** *Staphylococcus aureus*, enterotoxin, superantigen, SSL, MHC class II

## Abstract

*Staphylococcus aureus* has become a significant cause of health risks in humankind. Staphylococcal superantigens (SAgs) or enterotoxins are the key virulent factors that can exhibit acute diseases to severe life-threatening conditions. Recent literature reports *S. aureus* has steadily gained new enterotoxin genes over the past few decades. In spite of current knowledge of the established SAgs, several questions on these putative enterotoxins are still remaining unanswered. Keeping that in mind, this study sheds light on a putative enterotoxin SEl26 to characterize its structural and functional properties. *In-silico* analyses indicate its close relation with the conventional SAgs, especially the zinc-binding SAgs. Additionally, important residues have been predicted that are vital for T-Cell receptor (TcR) and major histocompatibility complex class II (MHC-II) interaction and compared them with established SAgs. Besides, our biochemical analyses exhibited the binding of this putative enterotoxin with MHC-II, followed by regulating pro-inflammatory and anti-inflammatory cytokines.

## Introduction

*Staphylococcus aureus* is an opportunistic pathogen that causes a wide range of infections, ranging from mild skin infections to life-threatening conditions. One of the most fascinating aspects of *Staphylococcus aureus* is its ability to produce superantigens (SAgs), which play a crucial role in its pathogenesis. SAgs, single-chain staphylococcal enterotoxins (SEs) with a molecular weight ranging from 22-29 kDa have been reported to cause transient food poisoning, skin infections, Kawasaki syndrome, arthritis, rheumatic fever and toxic shock syndrome (Argudín *et al.,* 2010; Yeung *et al.,* 2004; Gerlach *et al.,* 2017; Krakauer *et al.,* 2016). Unlike traditional antigens that activate a limited number of T-cells through specific interactions with the T-cell receptor (TcR) and major histocompatibility complex class II (MHC-II) molecules, SAgs have the ability to stimulate a large number of T-cell subpopulations (∼20%) in a non-specific manner followed by cytokine surge in the host (Scherer *et al.,* 1993; Li *et al.,* 1999; Sundberg et al., 2002).

Crystal structures of staphylococcal superantigens show that SAgs share a characteristic 3D structure comprising of an N-terminal OB-fold and a C-terminal β-grasp domain (Petersson *et al*., 2002; Fernández *et al*., 2006; Rödström *et al*., 2016). Although having common structural architecture, SAgs possess structural diversity in complex formation with MHC class II molecules. Previous studies showed that SAgs possess two independent binding sites for MHC-II: a low-affinity binding site (Kd ∼ 10^-5^ M) on the α-chain of MHC-II and a zinc-dependent high-affinity binding site (Kd ∼ 10^-7^ M) on the β chain of MHC-II (Fernández et al., 2006). Based on the sequence analysis and binding pattern studies, it was observed that some monovalent SAgs (SEB, SEC1-3, SEG, TSST1) could bind the MHC II α-chain through the low-affinity binding site present in the N-terminal OB domain, other monovalent SAgs (SEH, SEI, SEJ, SEK, SEL, SEM) can interact with the MHC-II β-subunit in a zinc-dependent manner through the high-affinity binding site present in the C-terminal β-grasp domain (Rödström et al., 2014; Sundberg et al., 2003; Petersson et al., 2001; Fernández et al., 2006). Moreover, bivalent SAgs (SEA, SED, SEE) can cross-link with two MHC-II molecules by concurrently binding the α-chain of one MHC-II and β-chain of other MHC-II molecule through both the low and high-affinity sites, respectively. Interestingly, bivalent SEA is found to be one of the most potent T-cell mitogenic SE (Abrahmsén *et al*., 1995). SEA interacts with MHC-II α using its N-terminal domain, which is also homologous to the MHC-II binding site of SEB (Jardetzky *et al*., 1994; Petersson *et al*., 2002). Besides, SEA cross-links with His81 of MHC-II β in a zinc-dependent manner (Abrahmsén *et al*., 1995; Hudson *et al*., 1995). Previously, SEA•(MHC-II)_2_ hetero-trimer was isolated, which suggested that SEA can simultaneously bind two MHC-II molecules expressed by the antigen-presenting cells (Tiedemann *et al*., 1995). Both the low-affinity and high-affinity sites are required for the superantigenic function of SEA. MHC-II bound SAg is highly requisite for maximum T-cell stimulation by SEA (Tiedemann *et al*., 1996).

Besides, a class of proteins called staphylococcal superantigen-like proteins (SSLs) contains 14 members and share structure-sequence similarity with conventional SAgs. Though there is no report so far regarding interactions of SSLs with TcR or MHC-II, its role in staphylococcal pathogenesis is important. SSLs lack mitogenic activity, despite having structural and sequence similarities to common superantigens (Arcus et al., 2002; Fraser & Proft, 2008). SSLs target the host immune proteins and carry out a variety of functions. SSL1 can interact with human ERK2; SSL5 and SSL11 bind P-selectin glycoprotein ligand 1 (PSGL-1), and SSL7 interacts with immunoglobulin IgA (Dutta *et al*., 2020; Bestebroer *et al*., 2007; Langley *et al*., 2005).

The staphylococcal enterotoxin gene family contains 24 genes (SEA-E, SEG-SEI, SElJ, SEK-SET, SElU-Y). In addition to these existing genes, a few more putative staphylococcal enterotoxin-like (SEl) proteins, namely SElZ, SEl26, SEl27, SEl28, SEs-2p, SEl29p, SEl30, SEl31, SEl32 and SEl33 have been identified but not characterized (Dicks *et al.,* 2021). Here, an attempt has been made to characterize SEl26 as a member of the enterotoxin family and to understand its MHC-II binding and cytokine regulation.

## Materials and methods

### Phylogenetic and sequence alignment analysis

BLASTp (Boratyn *et al*., 2012) identified the novel putative enterotoxin SAOUHSC_01705 in *S. aureus* NCTC8325 as a homologue of SEl26. SAOUHSC_01705 and other superantigens sequences were aligned by the ClustalW tool embedded in the MEGA11.0.11 software (Tamura *et al*., 2021). Neighbour-joining phylogenetic tree was constructed with 500 bootstrap replications. For sequence analysis, multiple sequence alignment was performed using Clustal Omega (Sievers *et al*., 2011). Protein function was assessed through structure or sequence similarities. The InterPro database (Paysan-Lafosse *et al*., 2023) was used to identify the domains in protein structures and sequences.

### Structure modelling of SAOUHSC_01705

Structure prediction of full-length SAOUHSC_01705 protein was performed in a standalone version of ΑlphaFold2 (Jumper *et al*., 2021), as implemented in ColabFold (Mirdita *et al*., 2022). Model prediction was run on a local computer with Ubuntu 18.04 operating system and accelerated with NVIDIA GeForce RTX 3060Ti GPU. Ligand bound SAOUHSC_01705 model was generated using AlphaFill (Hekkelman *et al*., 2023).

### Molecular docking of SAOUHSC_01705 with MHC-II

HADDOCK docking server (Dominguez *et al*., 2003) generates the complex models based on the FFT correlation algorithm and ranks the model according to their clustering properties, followed by the generation of four types of models based on balanced, hydrophobic, electrostatic and van der Waals plus electrostatic interactions. SAOUHSC_01705-apo and zinc-bound models were docked against the crystal structure of MHC-II using the HADDOCK 2.4.

### *In-silico* alanine-scanning mutagenesis

Computational alanine scanning mutagenesis was performed on six different protein complexes, SEA_D227A_•MHC-II (PDB: 1LO5; Petersson *et al*., 2002), SEB•MHC-II (PDB: 4C56; Rödström et al., 2014), SEC3•MHC-II (PDB: 1JWM; Sundberg *et al*., 2003), SEH•MHC-II (PDB: 1HXY; Petersson *et al.,*2001), SEI•MHC-II (PDB: 2G9H; Fernández *et al*., 2006) and docked structure of SAOUHSC_01705-MHC-II, using Robetta alanine scanning (Kortemme *et al.,* 2004). Briefly, two partners in each structure (the superantigen and the MHCII α or β) were selected, and all residues in the interface between the two partners were individually substituted to alanine. The predicted effect on the binding energy of the protein-protein complexes was computed to find the hot spots for complex formation. Hot-spot residues were defined as those for which alanine substitutions have predicted destabilizing effects on ΔΔG_complex_ of more than or equal to Rosetta energy units (approximating 1 kcal/mol).

### Recombinant protein production and purification

Staphylococcal SAOUHSC_01705 (UniProt: Q2FXX5) gene was amplified using the S.aureus NCTC8325 genomic DNA with the forward primer 5’-CGCGGATCCTTGTTAGAAGGAATTTTTAC-3’ and reverse primer 5’-CCCAAGCTTTTATGATTTGAATAAATAGATATC - 3’. Amplified PCR product was cloned into BamHI and HindIII digested pET28a expression vector. The recombinant plasmid containing SAOUHSC_01705 gene was transformed into E. coli BL21 (DE3) cells and further selected on kanamycin plates. Protein was affinity-purified using Ni-Ni-Sepharose High-Performance Affinity Matrix (GE Healthcare Biosciences) and eluted using 500mM imidazole containing buffer. Affinity-purified protein was further purified (>95%) using Superdex 75 prep-grade matrix in a 16/70 C column (GE Healthcare Biosciences).

### Custom antibody production and purification

Recombinant SAOUHSC_01705 protein (>95% purity) was administered to immunize rabbits. Rabbit maintenance, immunization and antiserum production was performed by Bio Bharati Life Science Pvt Ltd. Quality of SAOUHSC_01705 anti-sera was evaluated by ELISA. Polyclonal antibodies from antiserum were purified using Protein A Sepharose (Sigma, USA) following the manufacturer’s protocol. Antiserum was diluted 2-fold in dilution buffer containing 10 mM Tris-HCl (pH 7.4) and loaded onto the column. Specific antibodies were finally collected in an elution buffer containing 100 mM Glycine (pH 2.7).

### Immunofluorescence analysis

Immunofluorescence laser scanning confocal microscopy imaging was performed using polyclonal ani-SAOUHSC_01705-antibody raised in New Zealand breed Rabbit and MHC-II antibody. To that end, 10ng PMA-differentiated THP1 cells were incubated with 1µg protein for 2hrs. After incubation, cells were washed thoroughly (thrice) with 1X PBS to remove unbound proteins. Washed cells were further fixed with 4% paraformaldehyde for 15mins followed by incubation with anti-SAOUHSC-antibody and anti-MHC-II antibody for 6hrs. Primary antibody-incubated cells were immune-stained with Alexa-dye conjugated anti-rabbit (Alexa Fluor 488) and anti-mouse (Alexa Fluor 568) secondary antibodies. After thorough washing, secondary antibody-incubated cells were mounted with ProLong Gold Antifade (Thermo Scientific, USA) reagent and visualized using Olympus FV3000 Confocal Laser Scanning Microscope (Olympus, Japan).

### Real-time polymerase chain reaction (qPCR)

Total RNA from THP1 cells was isolated using the TRIzol reagent following the manufacturer’s protocol. In brief, macrophages were incubated with or without SAOUHSC_01705 for 0hr or 2hr or 4hr using 5µg/ml protein. Further, cells were washed thrice with complete RPMI media and unbound proteins were removed. After washing, cells were again washed with 1X PBS and total RNA was extracted. 1µg of total RNA was converted to cDNA for each sample using iScript cDNA Synthesis Kit (BioRad, USA) and stored at -80 °C for further use. qRT-PCR was performed BioRad CFX96 Thermal cycler using BioRad SSOAdvanced Universal SYBrGreen Supermix (BioRad, USA). Human TNF-alpha, IFN-gamma, IL-10, iNOS and NF-kB mRNA levels were monitored for above mention samples. β-actin was used as the internal reference throughout the experiments. Livak method (2^-ΔΔCq^) was used to calculate the fold change in mRNA expression for individual samples.

## Results

### Phylogenetics and sequence analysis reveal SAOUHSC_01705 as an enterotoxin

InterPro search results reported that SAOUHSC_01705 contains an N-terminal OB-fold (InterPro entry IPR006173) and a C-terminal β-grasp domain (InterPro entry IPR006123) that matches with the established SAgs. BLASTp analysis of SAOUHSC_01705 (234 aa) showed a close relationship with staphylococcal superantigens whereas the putative enterotoxin is having distant relationship with SSLs (Figure 1a). Phylogenetic analysis further exhibited that SAOUHSC_01705 is more associated with the zinc-binding SAgs. In addition, SAOUHSC_01705 showed 30-40% sequence identities with the superantigens that possess zinc-dependent MHC-II binding regions (SEA: 41.7%, SEE: 40.4%, SED: 37.9%, SEH: 35.9%, SEI: 34.9%). Sequence alignment of SAOUHSC_01705 showed zinc binding conserved sequences (H-L/I/V/F-D region) in the C-terminal β-grasp domain (Figure 1b). These analyses indicated that SAOUHSC_01705 could bind zinc and interact with MHC-II in a zinc-dependent manner.

**Figure 1.**
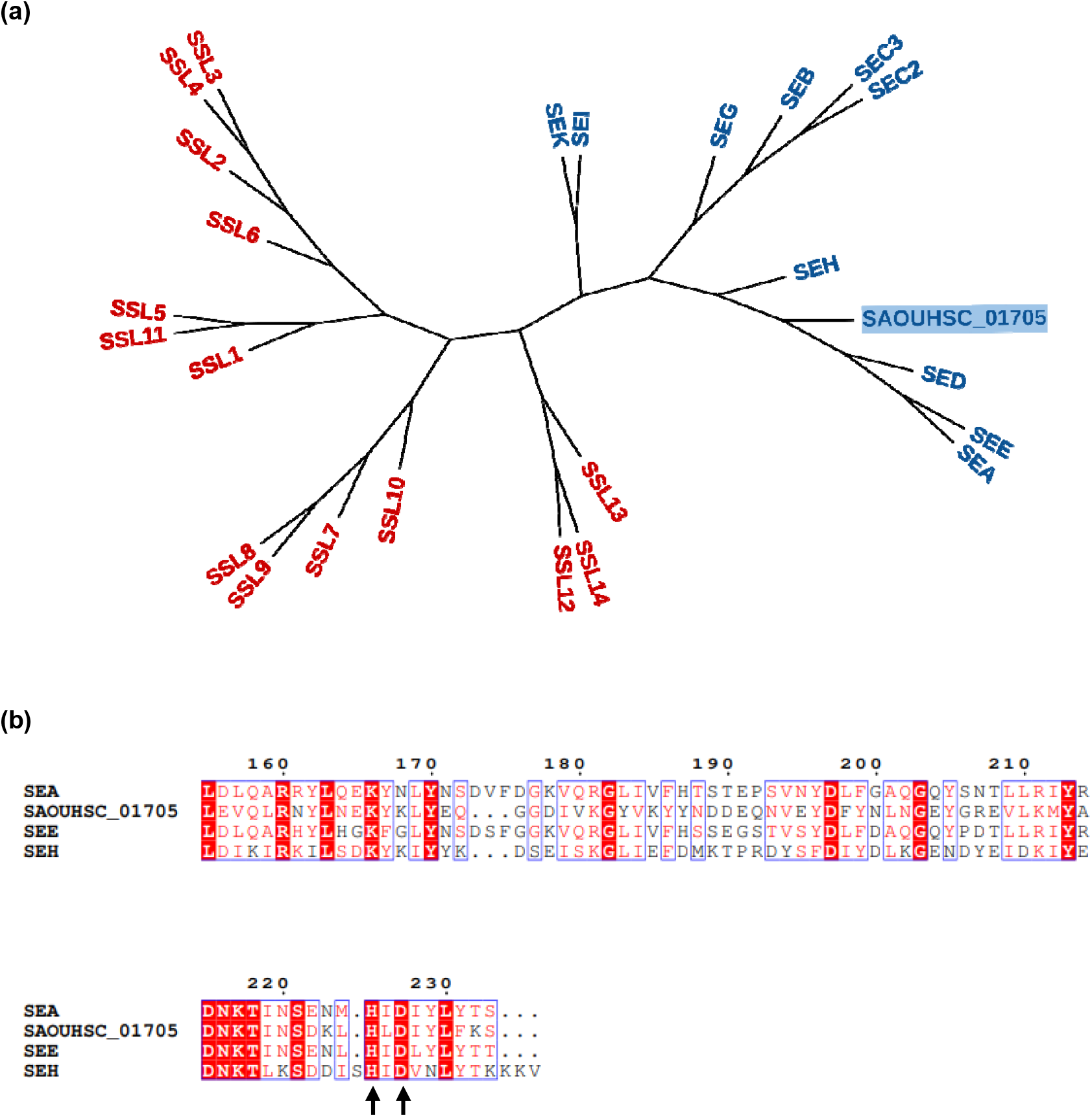
Phylogenetics and sequence analysis reveal SAOUHSC_01705 as an enterotoxin family member. (a) Phylogenetic tree analysis showing the relation of SAOUHSC_01705 with SAgs and SSLs. SAOUHSC_01705 possesses a close relationship with staphylococcal superantigens whereas this putative enterotoxin is having distant relationship with SSLs (b) Arrows indicating the histidine and aspartate residues in the c-terminal β-grasp domain involved in the zinc-binding which further interacts with MHC-II β subunit in the crystal structures of superantigens. Sequence alignment was performed with Clustal_omega, and the figures were generated using ESPript.

### The general architecture of SAOUHSC_01705 matches with conventional SAgs

The model structure showed that the SAOUHSC_01705 protein has a closely packed domain arrangement similar to other staphylococcal superantigen structures. The N-terminal OB domain (residues 38-120) comprises of a β-barrel structure, and the C-terminal domain (residues 121-234) is majorly constituted with a β-grasp motif (Figure 2a). In general, zn-independent superantigen interactions occur with MHC-II α-subunit and N-terminal OB-fold of superantigen. In contrary, zn-dependent interactions are found between MHC-II β and the C-terminal β-grasp domain. The β-barrel in the OB domain comprises of two β-sheets and is capped by an α-helix. Besides, β-sheet packing against a single α-helix is present in the C-terminal β-grasp motif. The model structure possesses close similarity to other staphylococcal superantigen crystal structures, with an overall root-mean-square-deviation of 0.84 Å to SEA, 1.04 Å to SEB, 1.21 Å to SEC3, 0.71 Å to SEE, 0.93 Å to SEH and 1.10 Å to SEI. Additionally, the model structure shows a distant structural relationship to the staphylococcal superantigen-like proteins, with an overall root-mean-square-deviation of 4.75 Å to SSL1, 8.81 Å to SSL3, 5.28 Å to SSL4, 4.09 Å to SSL5, 5.01 Å to SSL6, 11.41 Å to SSL7, 9.79 Å to SSL10 and 5.20 Å to SSL11. ΑlphaFill predicted that zinc could possibly bind at the H-L/I/V/F-D region of the C-terminal β-grasp domain. The zinc-bound model structure of SAOUHSC_01705 was obtained from Αlphafill (Figure 2b). Moreover, the zinc-binding site was observed to be present on the surface of the C-terminal β-sheet domain where His226 Nε2 and Asp228 Oδ2 coordinate with the zinc ligand (Figure 2c).

**Figure 2.**
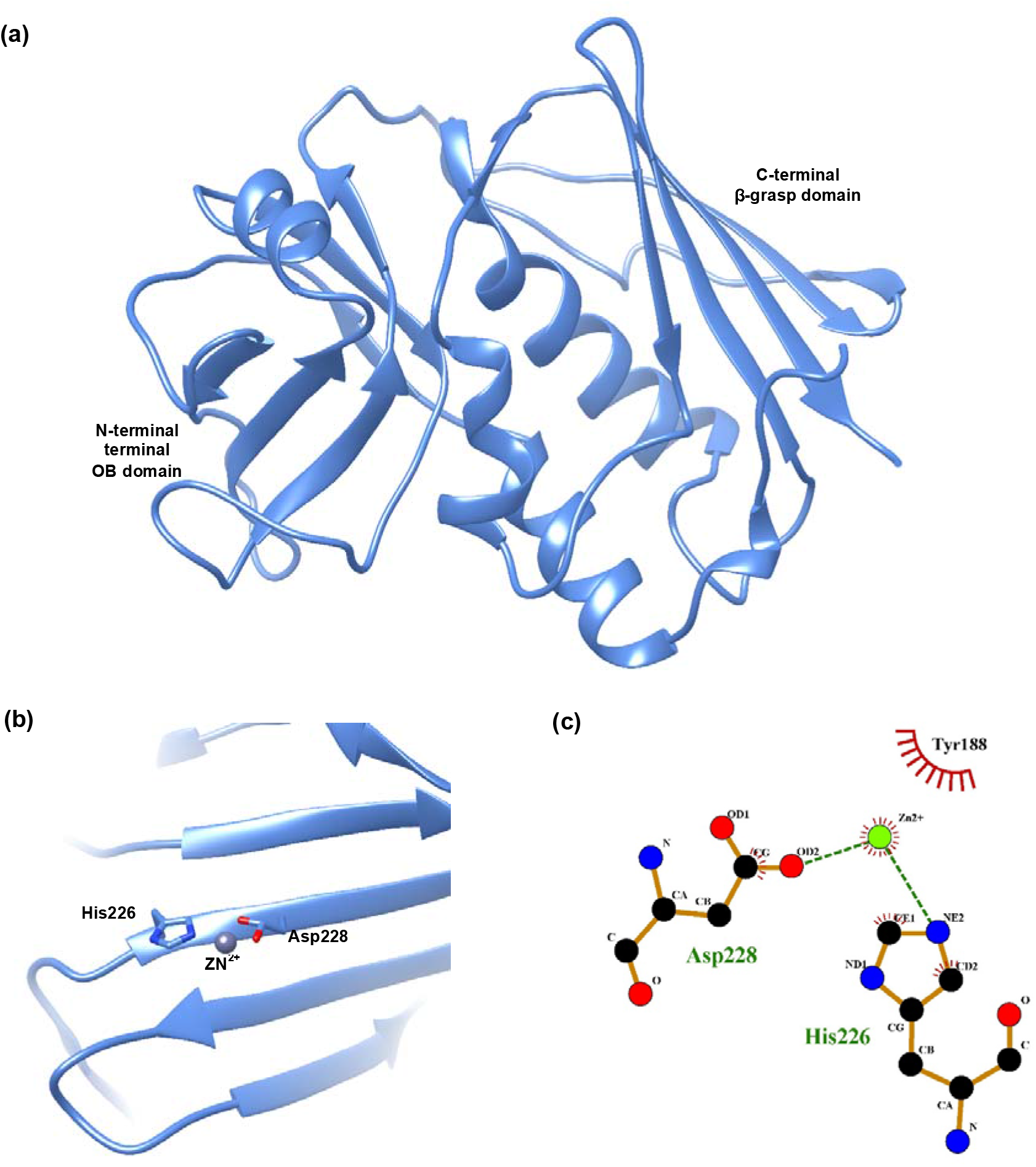
Predicted structural architecture of SAOUHSC_01705. (a) Ribbon representation of predicted SAOUHSC_01705 model, (b) Close view of zinc-binding region in the C-terminal beta-grasp domain, (c) Zinc coordination using His226 NE2 and Asp228 OD2.

### The interface of SAOUHSC_01705 and MHC-II subunits exhibit a bipartite interaction

Selected docked structure for zinc-independent MHC-II α interaction with SAOUHSC_01705 showed that the interaction occurs mainly through the loop regions (Thr47-Leu51 and Gly99-Asn106) (Figure 3a). These loop regions protrude into the cleft of side chains formed by the α1 helix and β-sheet of the MHC-II α-subunit. Hydrogen bond information between the critical residues is listed in Table 1. Asp49s and Arg50s made hydrogen bonds with Lys65α and Asn16α (residues belonging to the α-chain of MHC-II have ‘α’ as a suffix, residues belonging to the β-chain of HLA-DR1 have ‘β’ as a suffix, residues belonging to MHC-bound peptide have ‘p’ as a suffix, and residues belonging to enterotoxin have ‘s’ as a suffix). Two hydrogen bonds are found to be formed between Arg208s and Gln55α. In addition, Glu74s and His101s make hydrogen bonds with Lys37α and Gln55α, respectively. Interestingly, Thr105s interacts with HA-peptide, specifically with the Lys315p reporting the importance of peptide in SAOUHSC_01705•MHC-II α interaction.

**Figure 3.**
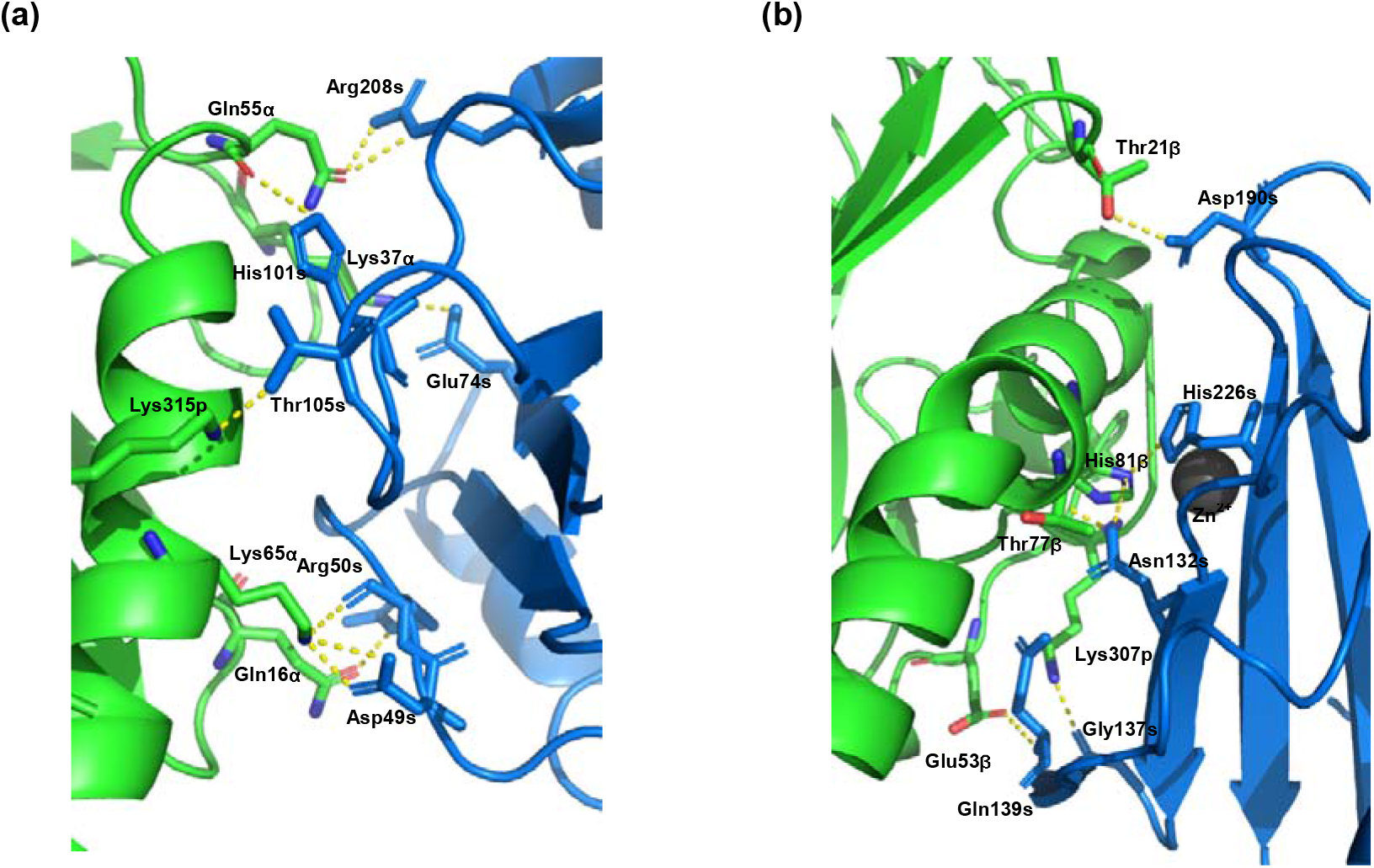
Ribbon diagrams of SAOUHSC_01705 MHC-II complex showing the interaction. SAOUHSC_01705 is blue, MHC-II chain is green and hydrogen bonds are represented as yellow dashed lines. (a) SAOUHSC_01705 is binding with MHC-II α chain using the loop region, (b) SAOUHSC_01705 is interacting with MHC-II β in a zinc-dependent manner.

**Table 1.** Interacting residues of SAOUHSC_01705 for MHC-II α and β cross-linking.

| SAOUHSC_01705•MHCII $\alpha$ interaction | | |
| --- | --- | --- |
| SAOUHSC_01705 residue | MHCII $\alpha$ residue | Distance (Å) |
| Asp49 | Lys65α | 2.60 |
| Asp49 | Lys65α | 3.10 |
| Arg50 | Asn16α | 2.70 |
| Arg50 | Lys65α | 2.80 |
| Glu74 | Lys37α | 2.70 |
| His101 | Gln55α | 3.30 |
| Thr105 | Lys315p | 2.80 |
| Arg208 | Gln55α | 2.70 |
| Arg208 | Gln55α | 3.10 |
| SAOUHSC_01705•MHCII $\beta$ interaction | | |
| Residue SAOUHSC_01705 | Residue MHCII $\beta$ | Distance (Å) |
| <u>Asn132</u> | Thr77β | 3.00 |
| <u>Asn132</u> | His81β | 3.30 |
| Gly137 | Lys307p | 2.70 |
| Glu139s | Glu53α | 3.50 |
| Asp190 | Thr21β | 2.70 |
| <u>His226</u> | His81β | 2.90 |

The final complex model for zinc-coordinated MHC-II β interaction was also selected from the balanced method. SAOUHSC_01705 binds with its concave β-sheet to the MHC-II molecule and HA-peptide. The total calculated contact region between SAOUHSC_01705 and MHC-II was found to be ∼1142 Å^2^. Interaction between enterotoxin and MHC-II was stabilized by zinc coordination. Residue His226, Asp228 from enterotoxin and His81 from MHC-II β-subunit coordinated with the zinc (Figure 3b). PDBePISA reported six hydrogen bonds to connect the SAOUHSC_01705 with MHC-II β (Table 1). Asn132s-Thr77β, Asn132s-His81β, Gly137s-Lys307p, Glu139s-Glu53α, Asp190s-Thr201β and His226s-His81β are the six hydrogen bonds that play a vital role in the interaction. Apart from the hydrogen bonds, several salt bridges formed between SAOUHSC_01705 and MHC-II also stabilized the interaction. Additionally, the HA-peptide governs part of SAOUHSC_01705 interaction on the MHC-II surface. The loop between Trp134s-Gln139s on SAOUHSC_01705 was found to be in close contact with the peptide. Gly137s formed hydrogen bond via its side chain oxygen with the nitrogen atom of Lys307p. Surprisingly, a hydrogen bond was noticed between Glu139s and Glu53α.

### Interaction of SAOUHSC_01705 with MHC-II is conserved with other SAg•MHC-II interactions

Upon analyzing the zinc-independent interaction, the total burying interface area of 598Å^2^ and 593 Å^2^ (total 1191 Å^2^) was observed for the SAOUHSC_01705 and MHC-II during the complex formation. However, in the case of SEA•MHC II α-interaction, the buried surface area was found to be 1121.5 Å^2^. Important residues for SAOUHSC_01705 in complex formation with MHC-II α-subunit are Asp49s, Arg50s, Glu74s, His101s, Thr105s and Arg208s. Other residues in the interacting region are Leu52s, His54s and Tyr96s. Structural superposition analysis showed that side chains of SAOUHSC_01705 in the loop region are lying with SEA. Arg50s of SAOUHSC_01705 was found to be conserved with SEA, and for Glu74, non-polar Asp70s is present in the corresponding position of SEA. Additionally, Leu52s, His54s and Tyr96s residues of SAOUHSC_01705 are highly conserved with SEA (Leu48s, His50s and Tyr96s; SEA-numbering). Overall, the interacting side-chain orientation is majorly similar in both SAOUHSC_01705 and SEA interaction with MHC-II α. In the case of zinc-mediated interaction, the docked structure of SAOUHSC_01705•MHC-II showed that zinc ion stabilizes the interaction by coordinating two ligands from enterotoxin SAOUHSC_01705 (His226 and Asp228) with His81 of MHC-II β subunit. The other vital residues Asn132s and Asp190s of SAOUHSC_01705, form hydrogen bonds with Thr77β and Thr201β of MHC-II β, respectively. Asn132s, His226s and Asp228s residues of SAOUHSC_01705 are conserved in staphylococcal SEH and SEI. The interface buries a dual surface area of 1247 Å^2^, with primary contributions from the β-subunit of MHC II. However, the buried surface of SEH and SEI are nearly the same, 1465 Å^2^ and 1461 Å^2^, respectively (Petersson *et al.,*2001; Fernández *et al.,*2006).

### Alanine scanning mutations on SAOUHSC reveal additional interacting residues for MHC-II interaction

Computational alanine scanning mutagenesis estimated the ΔΔG value of critical residues for complex formation between SAgs and MHC-II. Each interface residue was traded individually to alanine, and the substitution effect on the binding energy of the SAg•MHC-II complex was calculated as ΔΔG_complex_. Amino acids from SAgs having a destabilizing effect on the ΔΔG_complex_ of more than or equal to 1□kcal/mol after alanine substitution was considered to be hot-spot residues (Table 2). Most of the host-spot residues of SAOUHSC_01705 are familiar with SEH and SEI•MHC-II complex. The hot-spot residues of SEB and SEC3 were also analyzed for MHC-II interaction. In the case of SAg•MHC-II α subunit interaction, most of the hot-spot residues are common between SEA, SEB and SEC3 (Table 2) (Figure 5a). The analyses showed that Phe47 (SEA numbering, Phe44 in SEB and SEC3) is the most affected residue by alanine substitutions. Phe47 and Leu48 (SEA numbering) are also well-conserved in SEB and SEC3. However, there are five additional residues with a ΔΔG_complex_ value of more than 1 kcal/mol in the SEB and SEC3-MHC II complex, suggesting that the affinity may be higher than SEA•MHC II complex (Table 2). In the case of SAg•MHC-II β subunit interaction, two amino acids that were affected by alanine substitution in SAOUHSC_01705, SEH and SEI, are conserved amino acids. His226 in SAOUHSC_01705 and the corresponding His206s in SEH and His207s in SEI have the most pronounced effects (Table 2) (Figure 5b). Lastly, there are three additional residues with a ΔΔG_complex_ value of more than 1 kcal/mol in the SAOUHSC_01705•MHC II complex, suggesting that the affinity may be higher than SEH•MHC-II and SEI•MHC-II complex (Table 2). Interestingly, the amino acid with the most considerable ΔΔG_complex_ value (His226, SAOUHSC-numbering) has corresponding residues in SEH and SEI, which appear to be hot-spot residues for those complexes as well (Table 2). Asn132s and His226s (SAOUHSC_01705-numbering) residues are sequentially conserved among SAOUHSC_01705, SEH and SEI (Figure).

**Figure 4.**
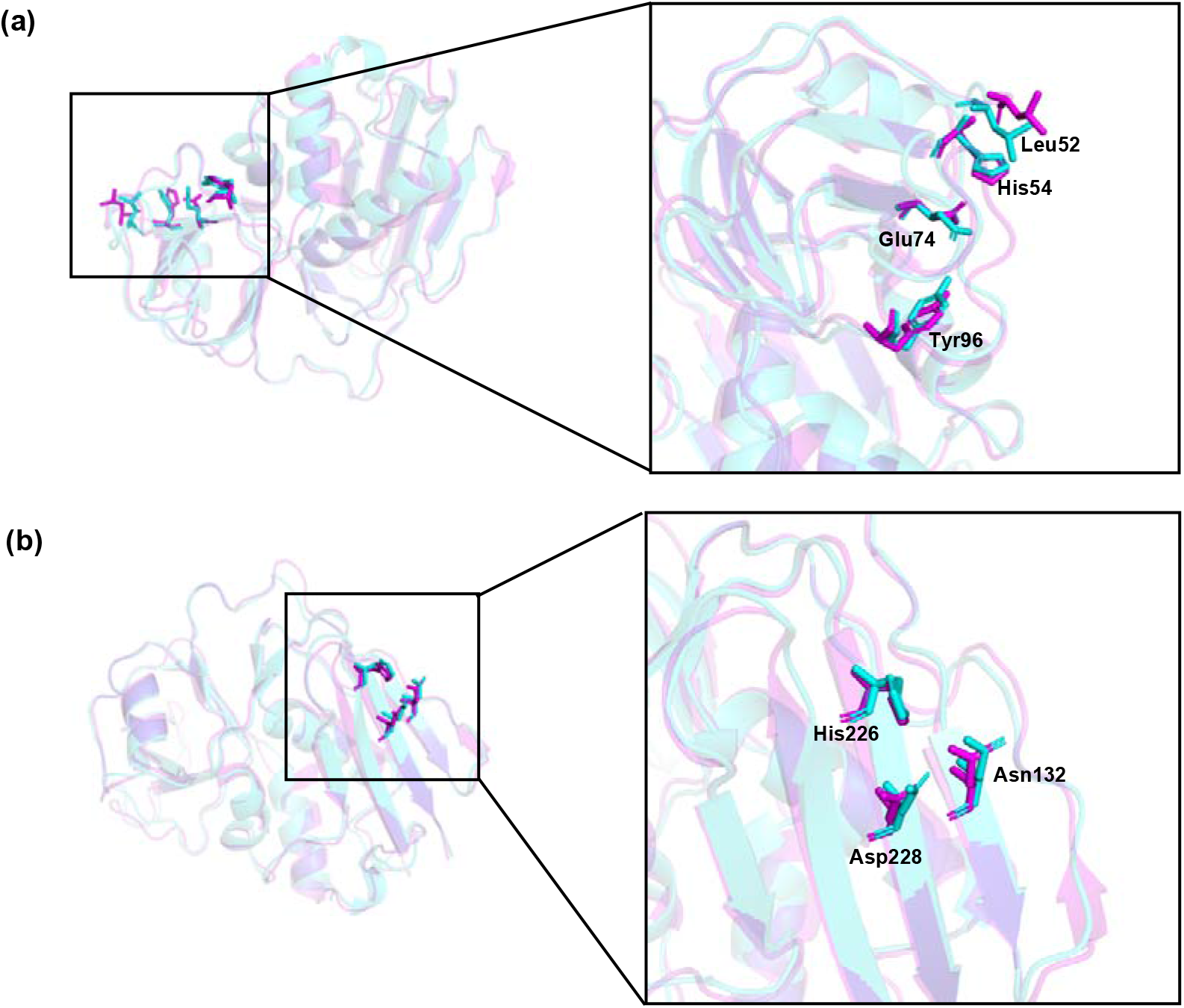
Conserved regions of SAOUHSC_01705 and SEA for MHC-II α and β interaction. Cyan is for SAOUHSC_01705, Pink is for SEA. (a) N-terminal OB domain conserved residues for MHC-II α binding (c) C-terminal β-grasp domain conserved residues for MHC-II β binding.

**Figure 5.**
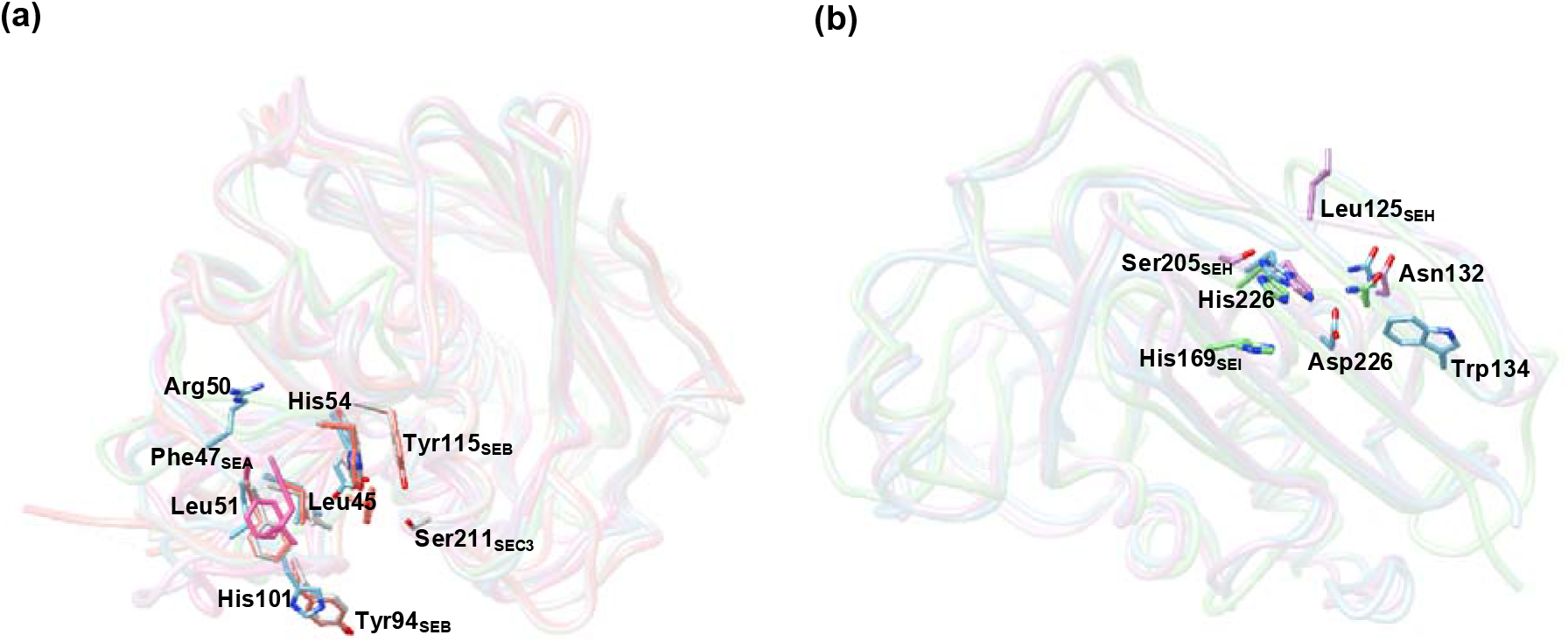
*In-silico* Alanine scanning mutagenesis showing the hot-spot residues in stick model. (a) SAOUHSC_01705 residues important for MHC-II α interaction, (b) SAOUHSC_01705residues important for MHC-II β interaction.

**Table 2.** Residues in the superantigens affected by *in-silico* alanine-scanning mutagenesis. ΔΔG_complex_ values are shown in kcal/mol. The conserved residues for MHC-II α or β interaction are represented in bold letters.

| Residue | $\Delta\Delta G_{\text{complex}}$ | Residue | $\Delta\Delta G_{\text{complex}}$ | Residue | $\Delta\Delta G_{\text{complex}}$ | Residue | $\Delta\Delta G_{\text{complex}}$ |
| --- | --- | --- | --- | --- | --- | --- | --- |
| SEA•MHCα | | SEC3•MHCα | | SEB•MHCα | | SAOUHSC•MHCα | |
| **Phe47** | 2.69 | **Phe44** | 2.52 | **Phe44** | 2.70 | Arg50 | 1.77 |
| **Leu48** | 1.89 | **Leu45** | 2.09 | **Leu45** | 2.01 | Leu51 | 1.09 |
|  |  | His47 | 1.62 | Phe47 | 1.63 | **Leu52** | 1.42 |
|  |  | **Glu67** | 2.09 | **Glu67** | 1.02 | His54 | 1.54 |
|  |  | **Tyr94** | 1.97 | Tyr89 | 1.32 | **Glu74** | 1.54 |
|  |  | **Tyr112** | 1.98 | **Tyr94** | 2.11 | His101 | 2.02 |
|  |  | Ser211 | 1.14 | **Tyr115** | 1.07 |  |  |
| SAOUHSC•MHCβ | | SEH•MHCβ | | SEI•MHCβ | | | |
| **Asn132** | 2.17 | **Asn113** | 1.16 | **Asn98** | 1.42 |  |  |
| Trp134 | 1.25 | Leu125 | 1.47 | His169 | 1.46 |  |  |
| **His226** | 2.93 | Ser205 | 1.49 | **His207** | 3.31 |  |  |
| Asp228 | 1.97 | **His206** | 1.61 |  |  |  |  |

### The overall model of TcR•SAOUHSC_01705•MHC-II quaternary complex

SAOUHSC_01705 also contains conserved T-cell receptor binding residues in its N-terminal OB-fold compared with SEA, SEB, SEC3 and SEE. Thus, it can be hypothesized that the putative SAOUHSC_01705 could be a potential superantigen that can bind both TcR and MHC-II in two different regions. Available TcR-bound crystal structures of SEA, SEB and SEC3 helped to construct a hypothetical TcR•SAOUHSC_01705•(MHC-II)_2_. SEA is the most toxic enterotoxin, where it mainly interacts with TcR using the α4 helix and β-turns from the β-barrel region of the N-terminal domain. SEA and SAOUHSC_01705 shared 41.7% sequence identity, where TcR and MHC-II-binding regions are highly conserved. The C_α_ alignment of SAOUHSC_01705 with SEA (RMSD 0.87 Å) indicated the high structural identity between the two structures and are more closely related. Staphylococcal SEA interacts with TcR engaging the shallow groove between the N- and C-terminal domains of the SAg. Residues from SEA that interacted with TcR were reported to be Asn25s, Tyr32s, Trp63s, Gly93s, Tyr94s, Val174s and Tyr205s (Rödström *et al.,* 2016). Surprisingly, all these residues except Tyr32s and Val174s are found to be conserved with SAOUHSC_01705, and side chains of these conserved residues are also lying in the orientation (Figure 6a). Upon complex modelling (PDBs used 1LO5 and 5FK9), it was observed that TcR lay in close proximity to the peptide binding groove of MHC-II. This information helped to model the TcR•SAOUHSC_01705•(MHC-II)_2_ structure (Figure 6b). The quaternary model showed that MHC-II interacts through the low-affinity binding site of SAOUHSC_01705 (present in the N-terminal domain) close to the TcR. In general, interaction between MHC-II and TcR in SAg-bound incidents produces more cytokines. Based on this, it can be suggested that SAOUHSC_01705 interacting with TcR and MHC-II (bivalently) can be a powerful enterotoxin as compared to others.

**Figure 6.**
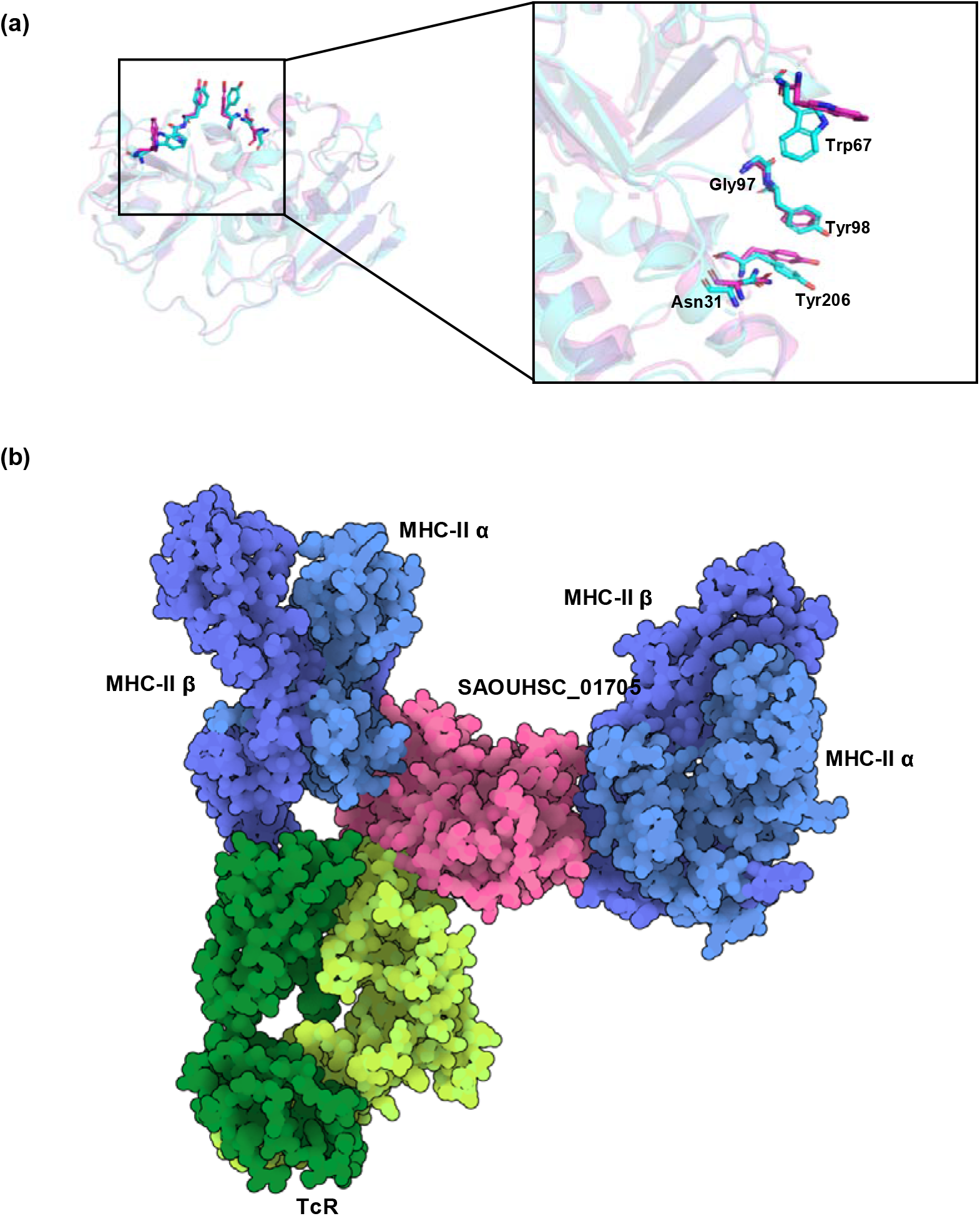
Conservancy between SEA and SAOUHSC_01705 and the hypothetical quaternary model of SAOUHSC_01705 in complex with TcR and two MHC-II molecules in Goodsell representation. (a) Important residues in SEA for TcR interaction is highly conserved in SAOUHSC_01705, (b) TcR alpha and beta chains are represented with light and dark green respectively. MHC-II alpha chain is represented with cyan and beta chain is represented with blue, SAOUHSC_01705 is represented with pink colour. An MHC-II molecule binds to the N-terminal domain of SAOUHSC_01705 and another MHC-II crosslinks to the C-terminal beta-grasp domain. Additionally, TcR interacts with SAOUHSC_01705 though the N-Terminal OB domain.

### SAOUHSC_01705 efficiently binds with macrophage-expressed MHC-II

The protein SEFT is a predicted superantigen. Since superantigens can directly bind to the MHC class II molecules outside the designated antigen binding domain, whether our predicted superantigen can also bind to the MHC II without any processing or not was verified by confocal microscopy. For this purpose, the THP-1 macrophages were incubated with purified recombinant SEFT followed by labelling with fluorescently-tagged anti-SEFT (Alexa fluor 488) and anti-MHC II (Alexa fluor 568) antibodies. BSA was used as a negative control in this experiment. In the SEFT–treated cells, the SEFT (green) and MHC II (red) were observed to be localized on the entire cell periphery. The yellow signal in the SEFT-treated cells, generated due to the overlapping green and red signal, indicates the co-localization of SEFT and MHC II on the macrophage plasma membrane (Figure 7). On the other hand, no such BSA localization or BSA/ MHC II co-localization was observed on the cell membranes of the control experiment. Therefore, the purified SEFT has the ability to bind the MHC II molecules without any antigen processing suggesting its superantigen-like property.

**Figure 7.**
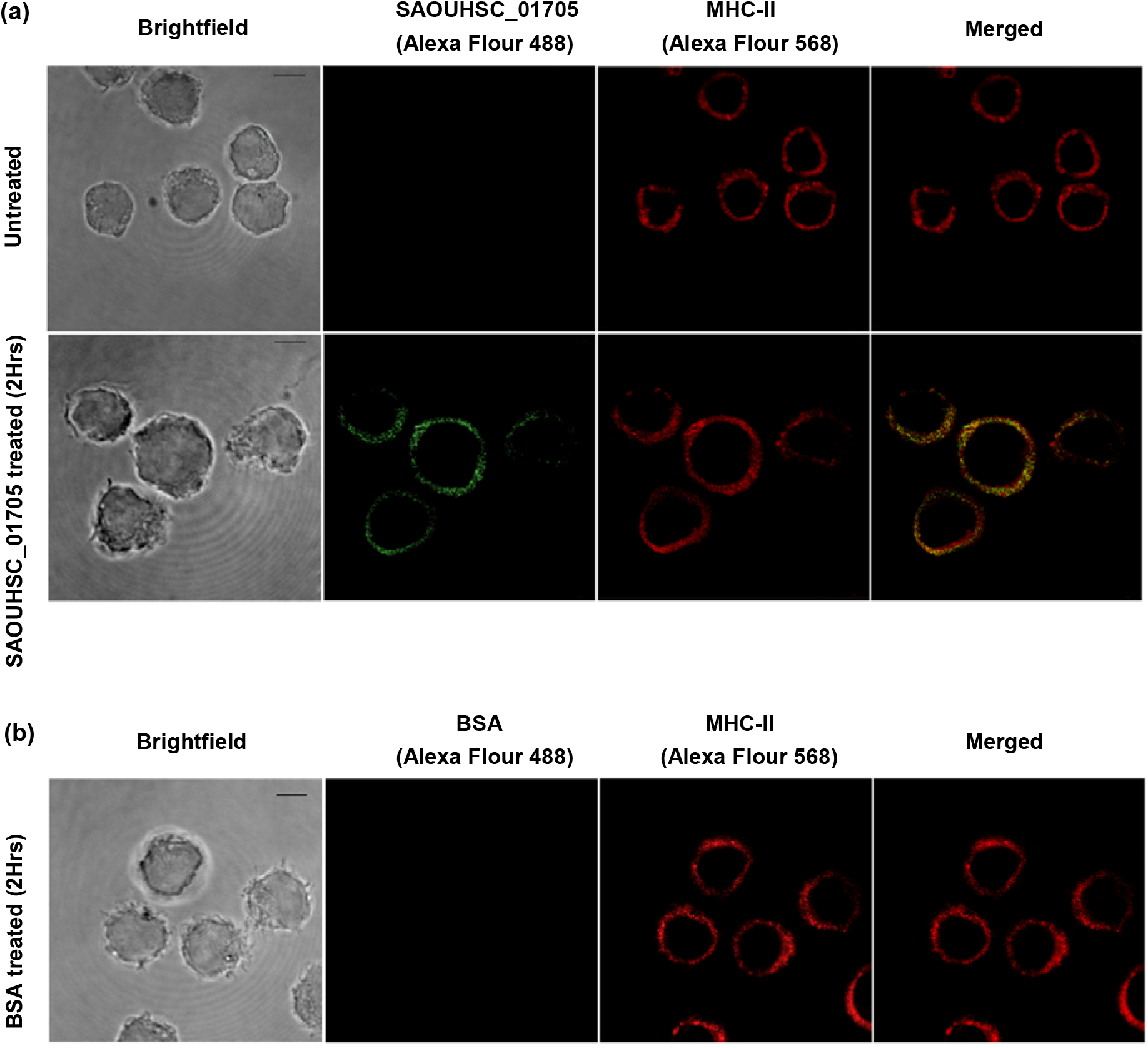
Confocal images confirm the cross-linking of SAOUHSC_01705 and BSA with MHC-II. (a) THP1 cells were incubated with anti-SAOUHSC_01705 antibody and anti-MHC-II antibody. No fluorescence was observed for Alexa Fluor 488 in case of untreated cells although fluorescence for anti-MHC-II antibody was observed. In case of treated cells, fluorescence for both anti-SAOUHSC_01705 and anti-MHC-II antibody was observed. Merged image panel exhibits the co-localization of toxin SAOUHSC_01705 and macrophage membrane expressed MHC-II, (b) BSA-treated cells do not exhibit fluorescence for Alexa Fluor 488. Also, merged image only contains fluorescence from Alexa Fluor 568 (anti-MHC-II antibody).

### MHC-II bound SAOUHSC_01705 can trigger anti-inflammatory responses

In order to understand the effect of SAOUHSC_01705 treatment in immune response patterns, alteration in expression of different cytokines, effector molecules and transcription factors under the treated condition compared to the untreated ones was checked at the transcript level. To accomplish this, the THP-1 macrophages were treated with the SAOUHSC_01705 for indicated periods (0 hr, 2 hr, 4 hr), followed by mRNA isolation and cDNA synthesis. The intracellular transcript level of all the cytokines was then analyzed by real-time PCR. Macrophage major cytokine TNFα level showed a significant and gradual decrease with time starting with a ∼1.5 fold decrease after 2 hrs and finally a ∼3.2 fold decrease after 4 hrs of treatment (Figure 8c). Another pro-inflammatory cytokine IFN-γ showed a similar trend where its expression was significantly reduced to ∼1.5 fold after 2 hrs and ∼2.5 fold after 4 hrs of treatment (Figure 8e). Since iNOS is crucial for effective bactericidal mechanisms, its intracellular level was also verified. Similar to the TNFα and IFN-γ, iNOS expression was also reduced to ∼2 fold and ∼3.5 fold after 2 hrs and 4 hrs of treatment, respectively (Figure 8a). Interestingly, the anti-inflammatory cytokine IL-10 showed the opposite expression pattern to that of pro-inflammatory cytokines. After 2 hrs of treatment, the IL-10 level significantly increased ∼3.75 fold, dropping to ∼2.2 fold after 4 hrs (Figure 8b). As lack of NF-κβ was reported to result in immune-compromisation, the intracellular mRNA level of NF-κβ was also checked. From the real-time RT-PCR, it was found that after 2 hrs of infection, the expression level of NF-κβ significantly increased ∼3 fold and then dropped to ∼1.5 fold after 4 hrs (Figure 8d).

**Figure 8.**
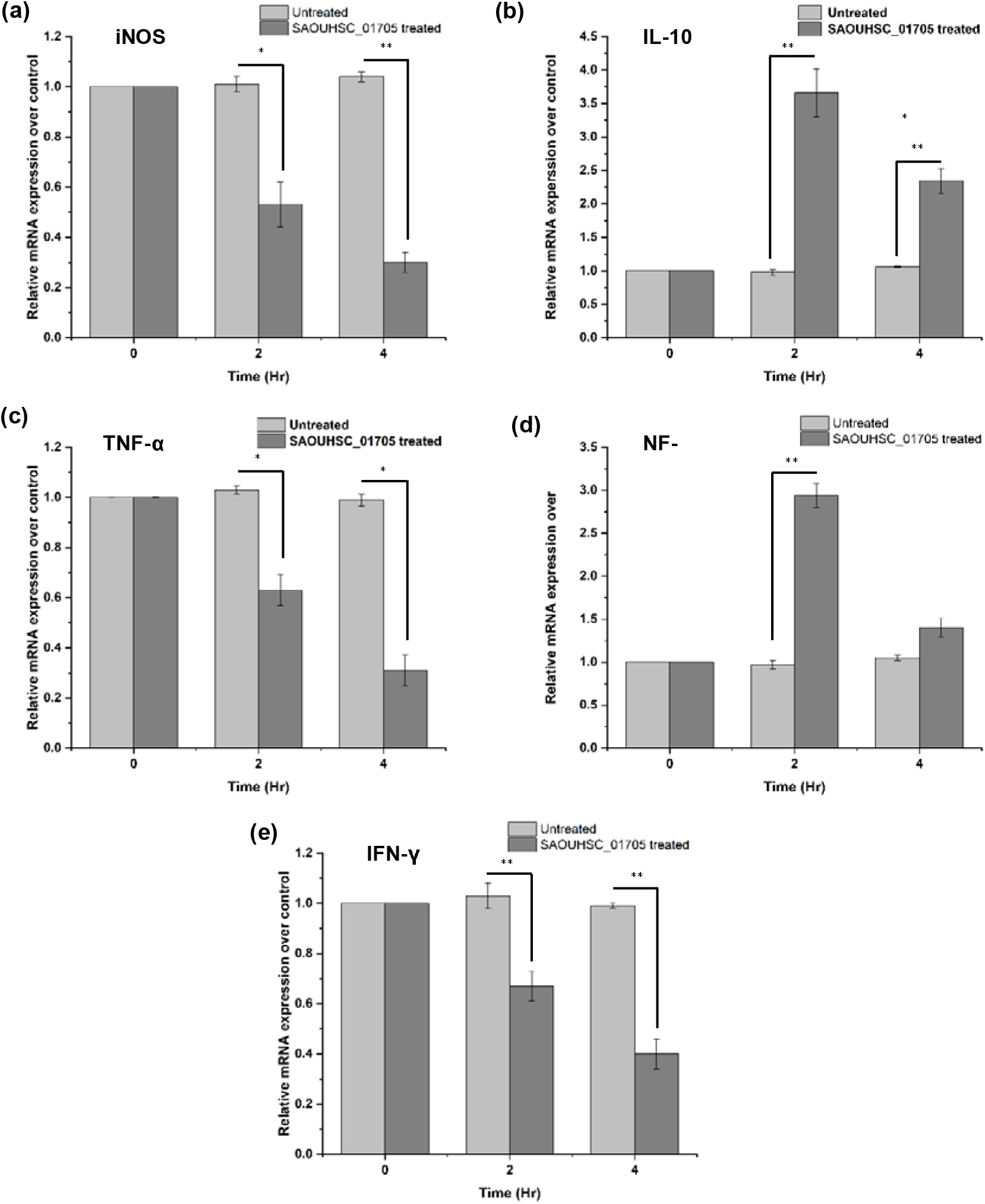
Relative mRNA expressions of pro-inflammatory and anti-inflammatory cytokines after SAOUHSC_01705 treatment. (a) iNOS expression was reduced to ∼2 fold and ∼3.5 fold after 2 hrs and 4 hrs of treatment, (b) IL-10 level significantly increased ∼3.75 fold after 2hrs of treatment then dropping to ∼2.2 fold after 4 hrs, (c) TNFα level showed a significant and gradual decrease with time starting with a ∼1.5 fold decrease after 2 hrs and finally a ∼3.2 fold decrease after 4 hrs of treatment, (d) the expression level of NF-κβ significantly increased ∼3 fold and then dropped to ∼1.5 fold after 4 hrs, (e) IFN-γ showed a similar trend where its expression was significantly reduced to ∼1.5 fold after 2 hrs and ∼2.5 fold after 4 hrs of treatment.

## Discussion

Staphylococcus aureus has become notorious due to its ability to emerge drug-resistance. *S. aureus* produces a collection of enterotoxins or superantigens that can activate a large number of T-cell subpopulations. MHC-II-SAg-TcR exhibits the most mitogenic effect, followed by the cytokine storm in the host body. These superantigens are one of the major virulent factors during pathogenesis. In this current study, a putative exotoxin enterotoxin family protein has been explored for its binding capability with MHC-II and T-cell receptors. The exotoxin was found to be closely related to the enterotoxin family protein, especially the zinc-binding superantigens. From the sequence information and modelled 3D, the zinc-binding region is identified. Besides, docking with MHC-II followed by molecular dynamics simulation has been performed to understand the interaction pattern. Alanine scanning mutagenesis of SAOUHSC_01705•MHC-II has been performed and compared with other SAgs. This study also investigates the conserved T-cell receptor binding region in the SAOUHSC_01705 toxin.

Phylogenetic analysis suggests putative SAOUHSC_01705 is a member of the staphylococcal enterotoxin family, and the toxin is not a part of staphylococcal superantigen-like proteins. Upon further analysis, it has been observed that SAOUHSC_01705 contains MHC-II α-binding conserved residues in its N-terminal domain. Previous reports show that SEA interacts with the α-chain of MHC-II using the N-terminal OB domain. Important residues for SEA•MHC-II α-interaction are reported to be Phe47, leu48, His50, Asp70 and Tyr92. Most of the critical residues, such as Leu52, His54 and Tyr96 of SAOUHSC_01705, are found to be highly conserved with SEA. It has been observed that corresponding conserved residues of SAOUHSC_01705 attend similar locations and side-chain orientations compared to SEA. Therefore, we predict that SAOUHSC_01705 can interact with MHC-II α through this conserved region. The enterotoxin also possesses a zinc-binding region in its C-terminal β-grasp domain. This zinc-binding region of SAOUHSC_01705 is highly conserved with other staphylococcal SAgs (SEA, SED, SEE, SEH, SEI). SEA uses His187, His225 and Asp227 to coordinate with zinc, SEH uses His206 and Asp208 for zinc coordination, and SEI coordinates with zinc through the residues His169, His207 and Asp209. MHC-II β binding to these zinc-binding SAgs occurs where His81 coordinates with SAg-bound zinc. Sequence analysis shows SAOUHSC_01705 possess His226 and Asp228 in its C-terminal β-grasp domain. Corresponding zinc-binding residues of SAOUHSC_01705 occupy similar locations and side chain orientations compared to conventional zinc-binding SAgs. Therefore, the presence of a conserved zinc-binding region in SAOUHSC_01705 suggests that it can also cross-link with MHC-II β and exhibit the mitogenic effect. Bivalent SEA shows the highest potential among all staphylococcal SAgs. Interestingly, SAOUHSC_01705 has been predicted to possess two MHC-II binding sites; therefore, it may also be bivalent.

The model structure of SAOUHSC_01705 comprises an N-terminal OB domain and a C-terminal β-grasp domain similar to the conventional superantigens found in nature. Two MHC-II binding regions are present in the opposite position to each other. The metal-bound model shows that zinc is coordinating with His226 Nε2, Asp228 Oδ1 and Oδ2 atoms from the β-grasp domain. Docked complex of SAOUHSC_01705•MHC-II indicates the bivalent binding patterns between enterotoxin and MHC-II. Binding with MHC-II α-subunit occurs when interacting loops of SAOUHSC_01705 protrude into the MHC-II. Structural superposition shows that side-chain orientations of conserved residues are nearly the same in the interacting loop regions of both SAOUHSC_01705 and SEA. Regarding zinc-mediated interaction, SAOUHSC_01705 cross-link with MHC-II β-subunit like other zinc-dependent SAgs such as SEH and SEI. Like conventional SAgs, SAOUHSC_01705 also interacts with MHC-II β using the concave β-sheet present in the C-terminal β-grasp domain of the enterotoxin. The previous report shows mitogenic activity is higher if MHC-II interacts with TcR when both are bound to SAgs (*et al.,* 20XX). Interestingly, it has been observed that MHC-II α binding region is in close proximity to the TcR binding site of SAOUHSC_01705. Conserved and additional residues for SAg•MHC-II interaction have been identified using the computational alanine scanning mutagenesis approach. This method identified ‘hot-spot’ residues from SAgs that are crucial for optimal MHC-II interaction.

Recombinant protein has been produced in the *E.coli* expression system. The protein is found to be monomeric when purified through size-exclusion chromatography. SAOUHSC_01705 is a ∼27kDa protein, and the size falls within the SAg size range (23-29 kDa). Protein-specific polyclonal antibodies have been raised in rabbits and purified for further experiments. The predicted interaction of SAOUHSC_01705 with MHC-II was experimentally evaluated. In-vitro experiments show SAOUHSC_01705 interacts with macrophage cell membrane-expressed MHC-II. It is well-known that superantigens can directly bind with the MHC class II molecules on the plasma membrane without being processed intracellularly for antigen presentation (Fernández et al., 2006). Our predicted enterotoxin SAOUHSC_01705 has also exhibited this characteristic feature as observed by the immunofluorescence confocal microscopy. THP1 macrophages incubated with purified recombinant SAOUHSC_01705 exhibited the co-localization of MHC II molecules (red) and SAOUHSC_01705 proteins (green) on the cell surface which appeared as a yellow ring for co-localization indicating their direct interaction. Superantigens or enterotoxins have been shown to be potent virulence factors which manipulate the macrophage inflammatory responses (Pidwill *et al*., 2021; Tuffs *et al*., 2022. Since we found the SAOUHSC_01705 to interact well with the MHC II molecules like other superantigens, it was further investigated whether this protein has any contribution on manipulating the macrophage inflammatory responses. Real time RT-PCR indicated that incubation with SAOUHSC_01705 can induce the overexpression of anti-inflammatory cytokine (IL-10) while at the same time downregulates the expression of pro-inflammatory cytokines (TNFα, IFNγ, iNOS). From this result it may be inferred that by attenuating the pro-inflammatory cytokine production, *S. aureus* brings the inflammatory responses of macrophage under its control and thereby ensures its survival in the hostile environment within the macrophage

## Conclusion

Staphylococcal superantigens are the hyper immunostimulatory molecules that directly cross-link to host MHC-II molecules and TcR, forming a trimolecular heteromeric complex. Superantigens are known to non-specifically activate a large number of immune cells (∼20% of all T-cells), followed by the production of pro-inflammatory cytokines such as tumour necrosis factor α (TNFα) and interleukin-2 (IL-2). In this current study, a putative exotoxin protein has been identified with possible TcR and MHC-II (bivalently) binding sites. The exotoxin is allied with the enterotoxin family, specifically the zinc-binding superantigens. Bivalent binding with MHC-II has been explored, and important residues for complex formation have been predicted. Studies of SAOUHSC_01705•MHC-II and other SAg•MHC-II complex report information on common motifs and additional residues of SAgs that are vital for interaction. Cellular studies report that SAOUHSC_01705 can directly bind the MHC-II, further regulating the mRNA level of both the pro-inflammatory and anti-inflammatory cytokines. This is the first report on putative SAOUHSC_01705 suggesting it as a bivalent enterotoxin and identifying a common set of interaction patterns in SAg•MHC-II complexes. This study will thus help design broad-spectrum antagonists to revoke SAg•MHC-II complex formation and impede the mitogenic activity caused by superantigens.

## Author contribution

SR conceptualized the study, performed *in-silico* analyses, recombinant protein production and wrote the draft manuscript. SS performed confocal microscopy, cytokine studies and interpreted respective data. SBDG and AKD supervised the work and improvised the final manuscript.

## Acknowledgements

SR acknowledges the Indian Institute of Technology Kharagpur for individual fellowship. The authors would like to acknowledge Debabrata Dutta and Rituparna Saha, IIT Kharagpur, for critical suggestions.

