## Supplementary for "Putative staphylococcal enterotoxin possesses two common structural motifs for MHC-II binding"

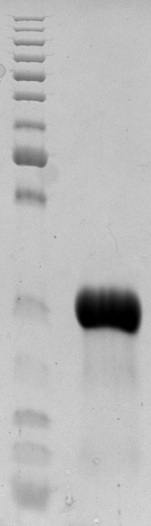

**27 kDa**

Supplementary Figure 1. Recombinant SAOUHSC_01705 toxin.

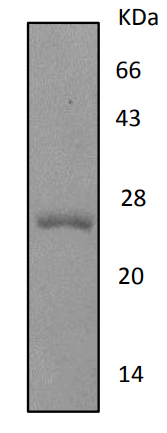

**27 kDa**

Supplementary Figure 2. Quality check of recombinant SAOUHSC_01705 for further antibody production.

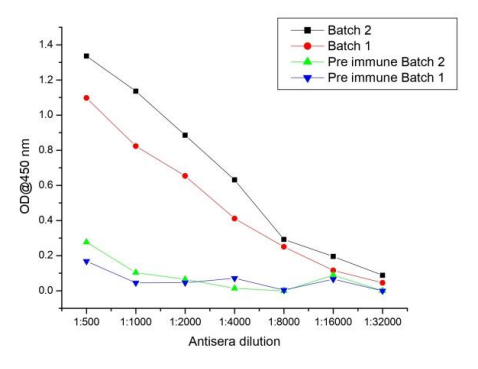
Supplementary Figure 3. Determination of titre of raised antisera containing anti-SAOUHSC_01705 antibody by ELISA.
